# A novel high throughput microwell outgrowth assay for HIV infected cells

**DOI:** 10.1101/2023.10.02.560422

**Authors:** Anthony D Fenton, Nancie Archin, Anne-Marie Turner, Sarah Joseph, Matthew Moeser, David M Margolis, Edward P Browne

**Author notes:** To whom correspondence should be addressed: Edward P Browne.

## Abstract

Although antiretroviral therapy (ART) is highly effective at suppressing HIV replication, a viral reservoir persists that can reseed infection if ART is interrupted. Curing HIV will require elimination or functional containment of this reservoir, but the size of the HIV reservoir is highly variable between individuals. To evaluate the overall size of the HIV reservoir, several assays have been developed, including PCR based assays for viral DNA, the Intact Proviral DNA Assay (IPDA), and the Quantitative Viral Outgrowth Assay (QVOA). QVOA is the gold standard assay for measuring inducible replication competent proviruses, but this assay is technically challenging and time consuming. To provide a more rapid and less laborious tool for quantifying cells infected with replication competent HIV, we developed the Microwell Outgrowth Assay (MOA), in which HIV infected CD4 T cells are cocultured with an HIV-detecting reporter cell line in a polydimethylsiloxane (PDMS)/polystyrene array of nanoliter sized wells (rafts). Transmission of HIV from infected cells to the reporter cell line induces fluorescent reporter protein expression that is detected by automated scanning across the array. We show that this assay can detect HIV infected cells with a high degree of sensitivity and precision. Using this approach, we were able to detect HIV infected cells from ART-naïve people with HIV (PWH) and from PWH on ART. Furthermore, we demonstrate that infected cells can be recovered from individual rafts and used to analyze the diversity of viral sequences. This assay may be a useful tool for quantifying and characterizing infected cells from PWH.

**Author summary:** Measuring the size of the HIV reservoir in people with HIV (PWH) will be important for determining the impact of HIV cure strategies. However, measuring this reservoir is challenging. We report a new method for quantifying HIV infected cells that involves culturing cells from PWH in an array of microwells with a cell line that detects HIV infection. We show that this approach can detect rare HIV infected cells and derive detailed virus sequence information for each infected cell.

## Introduction

Despite ongoing efforts, HIV remains an incurable infection leaving people with HIV (PWH) reliant on life-long regimens of antiretroviral therapy (ART) ^1^. While ART can prevent infection of CD4 T cells by HIV and subsequent HIV replication within these cells, a subset of cells will enter a state of latent infection ^2^. In these cells, HIV is reversibly silenced by a combination of factors, including low levels of key HIV activating transcriptional complexes and covalent modifications to proviral histones that promote heterochromatin formation ^3–5^. These latently infected cells evade clearance by the host immune system and can seed rebound viremia upon cessation of ART. As such, this reservoir represents the primary barrier to achieving an HIV cure^6,7^. Current HIV cure strategies aim to either induce permanent silencing of HIV or promote induction of viral antigen expression and clearance of infected cells ^4,6,8–13^. A handful of studies have examined the potential of latency reversal in both animal models and PWH and have shown some promise with regards to inducing expression of the reservoir but have thus far not been successful at reducing the size of the reservoir in PWH ^14–16^. Nevertheless, additional studies will be required to fully assess the potential of this approach. A key complication in terms of quantifying the outcome of a clinical intervention on the HIV reservoir is the complex nature of the reservoir. Of the residual integrated HIV proviruses in people of therapy, only a minor fraction are intact viruses – the remainder containing either large inactivating deletions or hypermutation ^17,18^. Furthermore, only a subset of the intact viruses reactivate expression in response to even “potent” latency reversing agents (LRAs)^19^.

To determine the effectiveness of any HIV cure strategy that targets the reservoir, a simple and scalable assay that measures replication competent and reactivation competent viruses would be useful. Thus far, several HIV reservoir assays have been developed. PCR based assays that measure the latent reservoir by quantifying HIV DNA have been widely used. However, PCR based methods that amplify HIV DNA tend to overestimate the intact reservoir due to the presence of defective proviruses ^20^. Droplet digital PCR (ddPCR) methods, such as the Intact Proviral DNA Assay (IPDA) attempt to combat this issue. The IPDA amplifies two HIV regions that are preferentially maintained in intact viruses and quantifies proviruses that are ddPCR positive for both. While this allows for a more accurate estimate of the intact reservoir size, some viruses with deletions or mutations in other regions of the HIV genome are still included, likely still leading to an overestimation of the intact reservoir size^20,21^. Additionally, polymorphism at the primer binding site can preclude detection of proviruses by the IPDA.

The quantitative viral outgrowth assay (QVOA) has been the gold standard for HIV cure clinical studies thus far ^22,23^. The QVOA uses limiting dilution, viral outgrowth within CD4 T cells and wells are assessed for p24 production by Enzyme-Linked Immunosorbent Assay (ELISA) and binarily scored. To quantify the size of the reactivation competent/replication competent reservoir, maximum likelihood statistics are then applied to the data to calculate the infectious units per million cells (IUPM). While this technique ensures that the estimate only includes replication competent virus, even with maximum stimulation of CD4 T cells from aviremic participants, in many cases, not all proviruses are reactivated with one round of stimulation^24^ . This means that the QVOA can underestimate the true size of the viral reservoir. Additionally, the QVOA is technically challenging and requires roughly 14-19 days to complete due the need to culture cells long enough to allow sufficient virus expansion for detection via p24 ELISA. These features limit the scalability of the assay as well as throughput, making its deployment in large clinical trials challenging.

To address this need, we developed a cell-based assay that uses novel CellRaft technology in combination with a highly sensitive Tat-dependent mCherry reporter cell line to detect HIV infection events, enabling rapid and sensitive quantification of HIV outgrowth from reactivated CD4 T cells from PWH. CellRafts are nanoliter sized wells (herein referred to as rafts) arranged in polydimethylsiloxane (PDMS)/polystyrene microarrays that allow single cell resolution microscopy in addition to extraction of individual rafts for sequencing purposes ^25,26^. Automated brightfield and fluorescence microscopy scanning of the arrays is enabled by a machine called the AIR system. The CellRaft array provides an environment where reporter cells are segregated from one another by the walls of each individual raft, allowing for analysis of rare events such as HIV viral reactivation while limiting spread of the virus into neighboring rafts. These events are quantified by automated scanning and image analysis and ultimately used to calculate an infectious unit per million (IUPM) based on the number of donor cells seeded on the array, reducing the physical labor required for the assay.

## Results

### CellRaft microarrays allow imaging and extraction of individual cells

CellRaft microarray technology enables the analysis of individual cells out of a bulk population via separation by PDMS walls. These arrays enable longitudinal culture of cells within the array and multiple arrays can be incorporated into the volume of a 96- well plate. The compact, biologically compatible design of the materials makes this technology useful for sensitive cell-based assays. In addition, the transparent nature of the devices allows for both brightfield and fluorescence microscopy directly within the array. Individual rafts can then be removed while target cells are still adhered to the raft. This allows extraction of rafts of interest within the array for other biochemical techniques such as PCR analysis of genes of interest (**Figure 1**). Critically, these processes are automated using the AIR system, an instrument and software suite that enables image capture and analysis, instrument control and extraction of individual rafts. Thus, we reasoned that co-culture of HIV infected cells with a HIV-activated fluorescent reporter cell line in CellRaft arrays might permit a more rapid and sensitive analog of the QVOA. We herein refer to this assay concept as the Microwell Outgrowth Assay (MOA).

**Figure 1:**
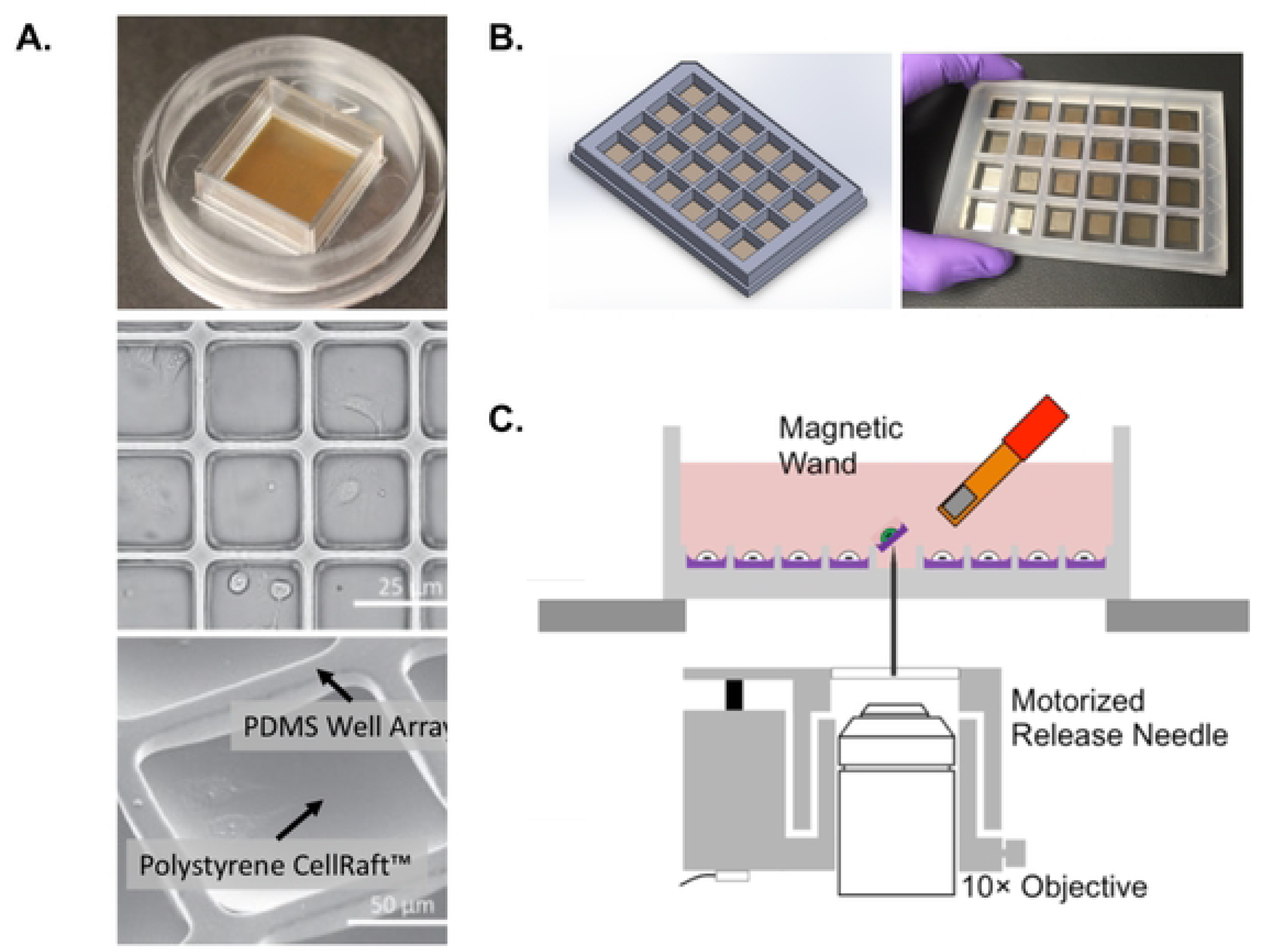
CellRaft arrays for the analysis of cell biology in extractable nanowells. **A.** Example of a 100μM CellRaft array with 40,000 magnetic polystyrene rafts surrounded by a polydimethylsiloxane (PDMS) scaffold. **B.** Example of a hexaquad array containing 154,000 rafts. **C.** Schematic of extraction technology for individual rafts. Individual rafts are ejected using a release needle and captured by a magnetic wand.

### Generation of a Tat-dependent reporter cell line to detect HIV infection

To achieve a novel, highly sensitive assay for quantification of latently infected cells in patient samples, we first generated a reporter cell line which expresses the fluorescent reporter protein mCherry in response to HIV infection. mCherry was selected due to its superior signal to noise ratio in CellRafts compared to green fluorescent protein (GFP). Human Osteosarcoma (HOS) cells were first transduced with a lentiviral construct containing an mCherry gene with expression controlled by an HIV long terminal repeat (LTR) and the HIV transactivation response element (TAR). An SV40 promoter driving expression of a puromycin resistance gene was included for selection of the resulting cells (**Figure 2A, 2B**). In this design, the TAR element from the reporter construct serves as a sensor of Tat expression during HIV infection and activates mCherry expression. The bulk puromycin selected cell population (HOS-LTR- mCherry) was then examined for the presence of HIV-responsive cells by infection with an HIV-GFP reporter virus that had been pseudotyped with the VSV-G envelope protein. Infection with the HIV-GFP virus resulted in a population of cells that was positive for both GFP and mCherry signal and an overall increased population of mCherry+ cells, indicating successful HIV infection coupled with activation of the mCherry reporter. Uninfected control HOS cells showed no mCherry fluorescence and uninfected HOS-LTR-mCherry cells showed significantly less mCherry activity suggesting the presence of clones with selective activation of the reporter in response to HIV infection (**Figure 2C**). Individual Tat-responsive clones within the puromycin resistant population were then enriched by transient transfection with Tat mRNA after plating on a CellRaft array, allowing us to rapidly screen the clones based on an increase in mCherry expression. Responsive clones were extracted from the array and expanded. We then further screened each clone by infection with VSV-G pseudotyped HIV-GFP, followed by flow cytometry. A clone with a strong mCherry response to HIV infection (A5) was selected for continued engineering (**Figure 2D**). To accommodate infection by clinical HIV isolates, the A5 clone was transduced with expression constructs for the CD4, CXCR4 and CCR5 receptors necessary for HIV infection, and stable expression was confirmed by flow cytometry (**Figure 2E**). The resulting cell line is herein referred to as MOA cells.

**Figure 2:**
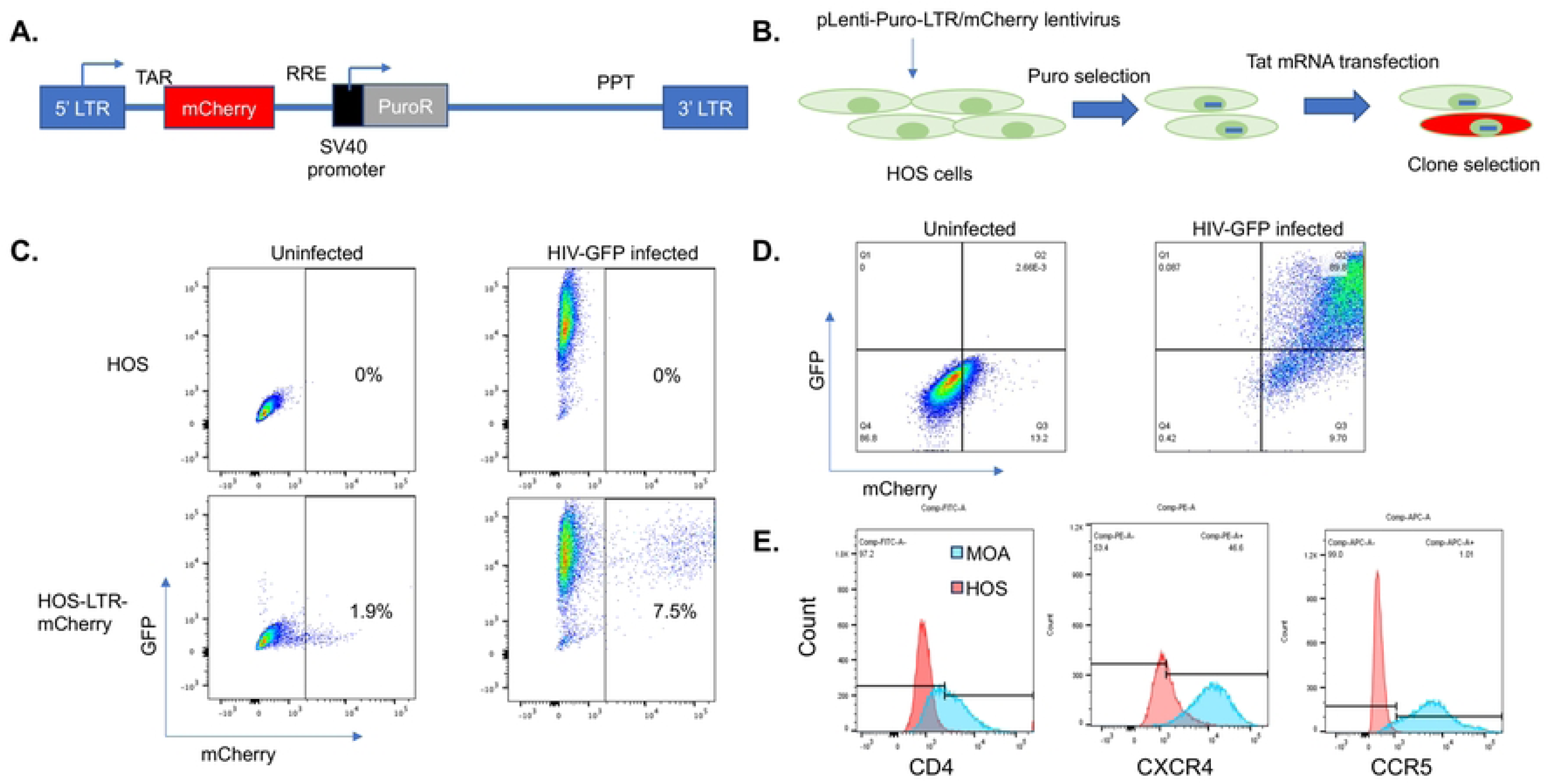
Generation of an HIV sensing cell line. **A.** Design of HIV reporter construct. An mCherry gene was cloned downstream of an HIV long terminal repeat (LTR) and HIV transactivation response element (TAR). Downstream of mCherry, a Rev response element (RRE) and internal SV40 promoter driving a puromycin resistance gene was added. A polypyrimidine tract (PPT) and a 3’ LTR were also included to allow generation of a packageable lentiviral RNA. **B.** Overall scheme for generation of an HIV sensing cell line shown. Human osteosarcoma (HOS) cells were transduced with lentiviral particles containing the construct from A and selected with puromycin. Cells containing an HIV responsive insert were enriched by transfection of Tat mRNA, and sorting mCherry+ cells using an AIR cell sorter. **C.** Flow cytometry of puromycin resistant HOS cell population in the presence and absence of infection with an HIV-GFP reporter virus. **D.** Flow cytometry showing the infection of a highly responsive HOS cell clone (A5) with HIV-GFP. **E.** Flow cytometry showing stable expression of HIV receptors CD4, CXCR4 and CCR5 in the HIV responsive A5 clone.

### Quantification of HIV infection in CellRaft arrays

To determine the ability of the MOA cells to detect and quantify replication competent HIV infection in the context of CellRaft arrays, MOA cells were plated in CellRaft arrays followed by infection of each array with varying volumes of the CXCR4- tropic HIV strain NL4-3. The arrays were then scanned each day for 5 days and the number of mCherry positive rafts were quantified for each timepoint (**Figure 3**). To select the optimal threshold of mCherry fluorescence to identify infected cells, we compared the number of “above threshold” rafts on infected and uninfected arrays at different thresholds and selected a threshold (5%) at which the false positive rate was below one in 10,000 rafts and the majority of true positive wells were detected (**S1 Fig**). We then tested different MOA cell plating densities and determined the optimal number of plated cells as four MOA cells per raft to achieve robust detection of infection events (**S2 Fig**). Also, we compared the signal from infected arrays cultured in different cell media (DMEM vs RPMI) and observed better performance in DMEM (**S3 Fig**).

**Figure 3:**
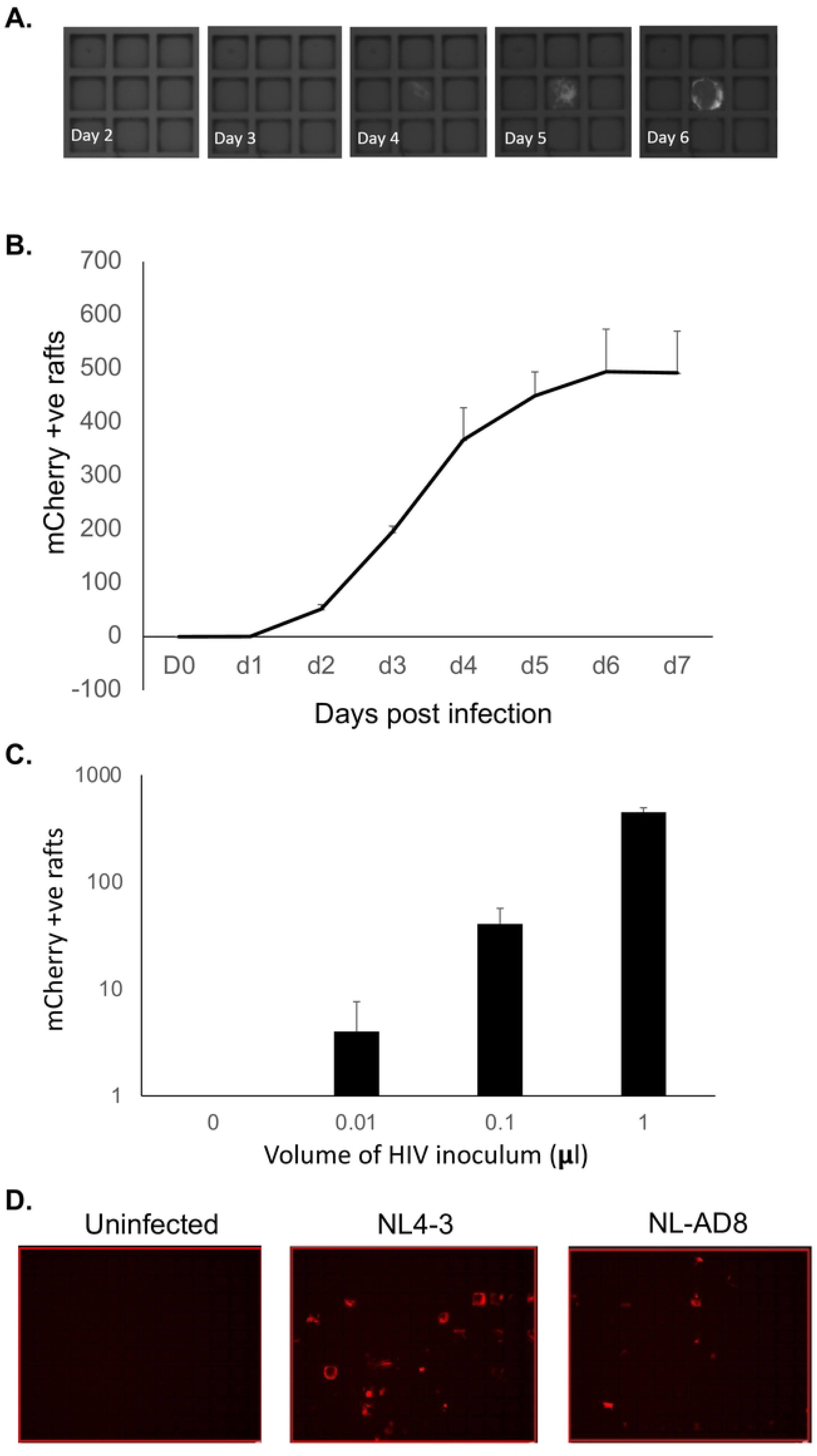
Detection of HIV infection in CellRaft arrays. MOA cells were plated in a CellRaft array, then infected with HIV (NL4-3) or mock infected. **A.** Representative images collected over time showing an individual positive raft. **B.** mCherry positive rafts were counted by scanning each day after infection of the arrays. The average number of positive rafts per array from three arrays is shown. Error bars represent standard deviation of the mean. **C.** At day 5 post infection, the number of mCherry positive rafts was determined for each volume of viral inoculum. The average number of positive rafts per array from three arrays is shown. Error bars represent standard deviation of the mean. **D.** MOA cells were plated in CellRaft arrays then viral inoculum from two different strains of HIV with different co-receptor tropisms (NL4-3, CXCR4 tropic and NL-AD8, CCR5 tropic) were added to the arrays. At 5dpi, the array was scanned for mCherry expression. Representative images are shown.

By examining the mCherry+ wells at each timepoint, we could observe a clear temporal pattern to the emergence of an infected raft signal. While the timing of mCherry expression varied from raft to raft, significant emergence of mCherry+ rafts was apparent by 2dpi, and increased rapidly until 4dpi, at which point the signal plateaued (**Figure 3A, 3B**). Additional mCherry+ rafts were not detected after 7dpi. When we compared the number of mCherry+ rafts at 5dpi across different concentrations of viral inoculum, there was a clear linear relationship between the volume of virus added and the number of mCherry+ rafts, suggesting that this approach can accurately and precisely measure small quantities of infectious HIV (**Figure 3C**). To confirm that these cells could also mediate infection by a CCR5 tropic strain, we next infected MOA cells in the CellRaft arrays in parallel with NL4-3 and the CCR5-tropic strain NL-AD8. We observed mCherry+ wells for both strains of HIV, indicating productive HIV infection by both CXCR4 and CCR5-tropic strains (**Figure 3D**).

### MOA detection of HIV infected CD4 T cells

We next examined the ability of MOA cells within CellRaft arrays to detect and quantify HIV infected CD4 T cells. CD4 T cells are the primary target of infection for HIV, and a large fraction of the latent reservoir resides within resting memory CD4 T cells in people with HIV (PWH). Thus, being able to detect infected CD4 T cells would be useful for measuring the abundance of infected cells in samples from PWH. To examine the ability of the MOA assay to detect and quantify HIV infected cells in CellRaft arrays, we first activated CD4 T cells from healthy donors, then infected them with a replication competent strain of HIV that encodes Heat Shock Antigen (HSA) in place of the Nef open reading frame. At 48h post infection we measured the abundance of productively infected CD4 T cells (HSA+) by flow cytometry (**Figure 4A**). Infected cells were then extensively washed to remove free virus particles and then different quantities of the infected cell suspension were cocultured with MOA cells in a CellRaft array for five days. We observed a linear quantitative relationship between the known abundance of infected cells plated in the array, and the number of mCherry+ rafts, suggesting accurate performance of the assay with respect to detecting and quantifying infected cells (**Figure 4B**). Furthermore, this assay exhibited a high degree of sensitivity, with as few as 1-2 infected cells per well of a quad array (6,400 rafts per well) being detected above background.

**Figure 4:**
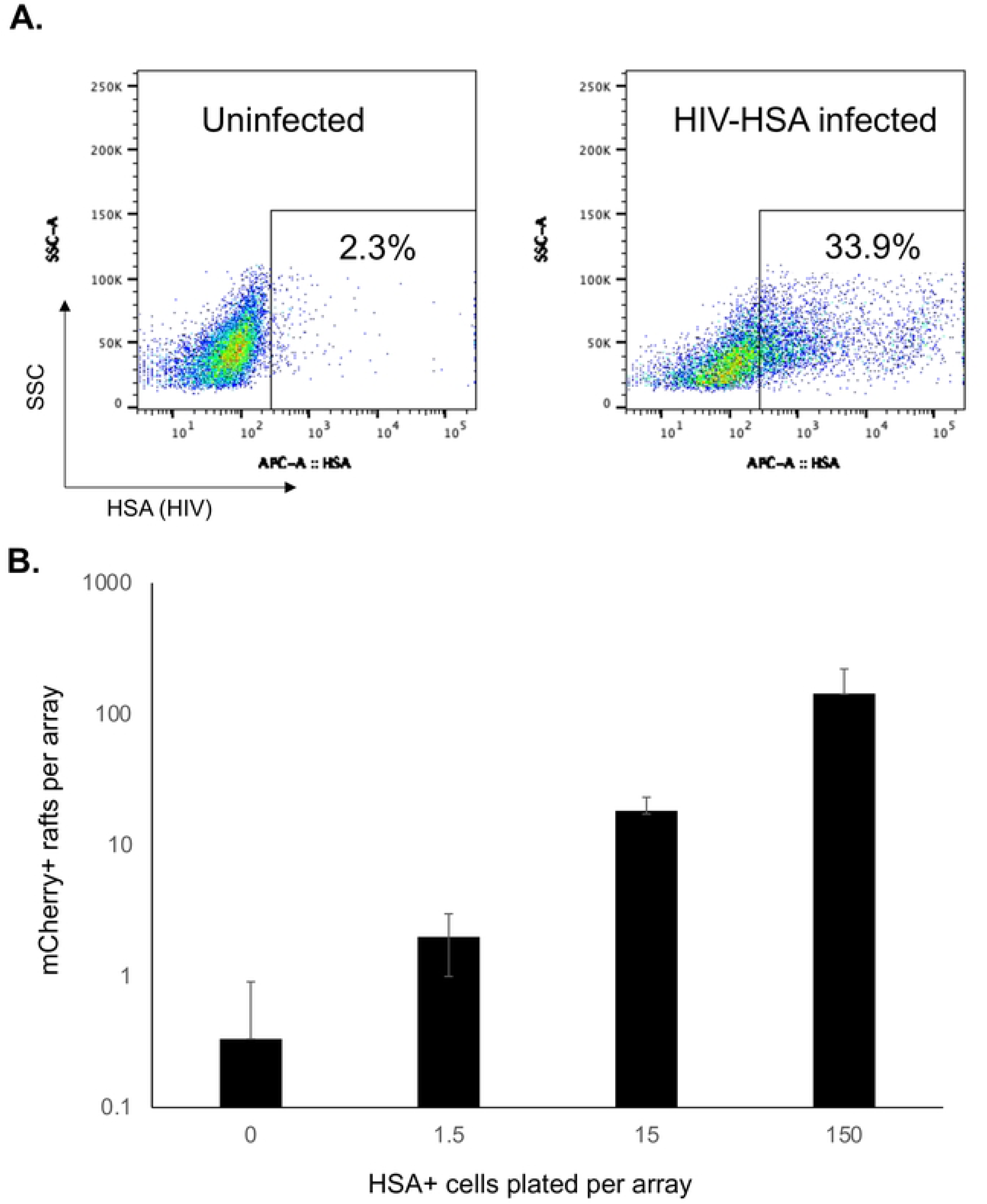
Detection of HIV infected CD4 T cells. **A.** Primary CD4 T cells from sero-negative donors were activated with PHA/IL-2 for 48h, then infected with a replication competent HIV strain that has been modified to encode Heat Shock Antigen (HSA) (HIV-HSA). At 48h post infection, the proportion of infected cells was determined by staining for HSA and flow cytometry. **B.** MOA cells were plated in CellRaft arrays and overlaid with varying amounts of cells from the infected culture. The arrays were scanned after five days of coculture and the number of mCherry positive rafts determined. Each bar represents the average of three replicate arrays. Error bars represent standard deviation of the mean.

### Optimizing MOA/CD4 T co-culture conditions for the microwell outgrowth assay

Detection of HIV infected cells from PWH will require coculture of high density CD4 T cells with MOA cells in the CellRaft arrays. To examine the capacity of the rafts to accommodate higher numbers of human CD4 T cells, we plated primary CD4 T cells at different densities per raft and observed that occupancy of the rafts saturated at ∼50 T cells per raft (**S4 Fig**). To induce viral gene expression and replication and transmission of HIV to the MOA cells, the CD4 T cells will also likely need to be activated prior to coculture. However, when we co-cultured MOA cells with high density PHA/IL-2 activated CD4 T cells we observed rapid killing of the MOA cells by the activated T cells (**Figure 5A**). This killing did not occur with resting CD4 T cells, demonstrating that activation of the T cells was required (not shown). Killing became apparent within 3-4 hours of coculture and resulted in almost complete loss of MOA cells by 24h in the presence of activated CD4 T cells. This phenomenon also occurred for CD4 T cells that were activated by CD3/CD28 stimulating beads, and by PMA/ionomycin, suggesting that this effect was not specific to PHA/IL-2 activation but was associated with general features of CD4 T cell activation (not shown). Viability of MOA cells was also not affected by supernatant from activated CD4 T cells, suggesting that direct contact between CD4 T cells and MOA cells was likely required for killing (not shown). Since killing likely depended on a signaling pathway that was induced during T cell activation, we tested a panel of inhibitors of T cell signaling pathways to determine whether they could block killing of MOA cells by the activated CD4 T cells. We observed that Dasatinib, a tyrosine kinase inhibitor, was able to rescue MOA cell viability in the presence of activated CD4 T cells, indicating that Dasatinib blocked a pathway that is essential for killing. By examining different Dasatinib concentrations, we further established that 10nM Dasatinib was sufficient to robustly preserve MOA cell viability during coculture (**Figure 5B**). Higher Dasatinib concentrations (1-10μM) still protected the MOA cells from CD4 T cells but caused a noticeable reduction in MOA cell growth suggesting possible cytotoxic or cytostatic activity in this concentration range.

**Figure 5:**
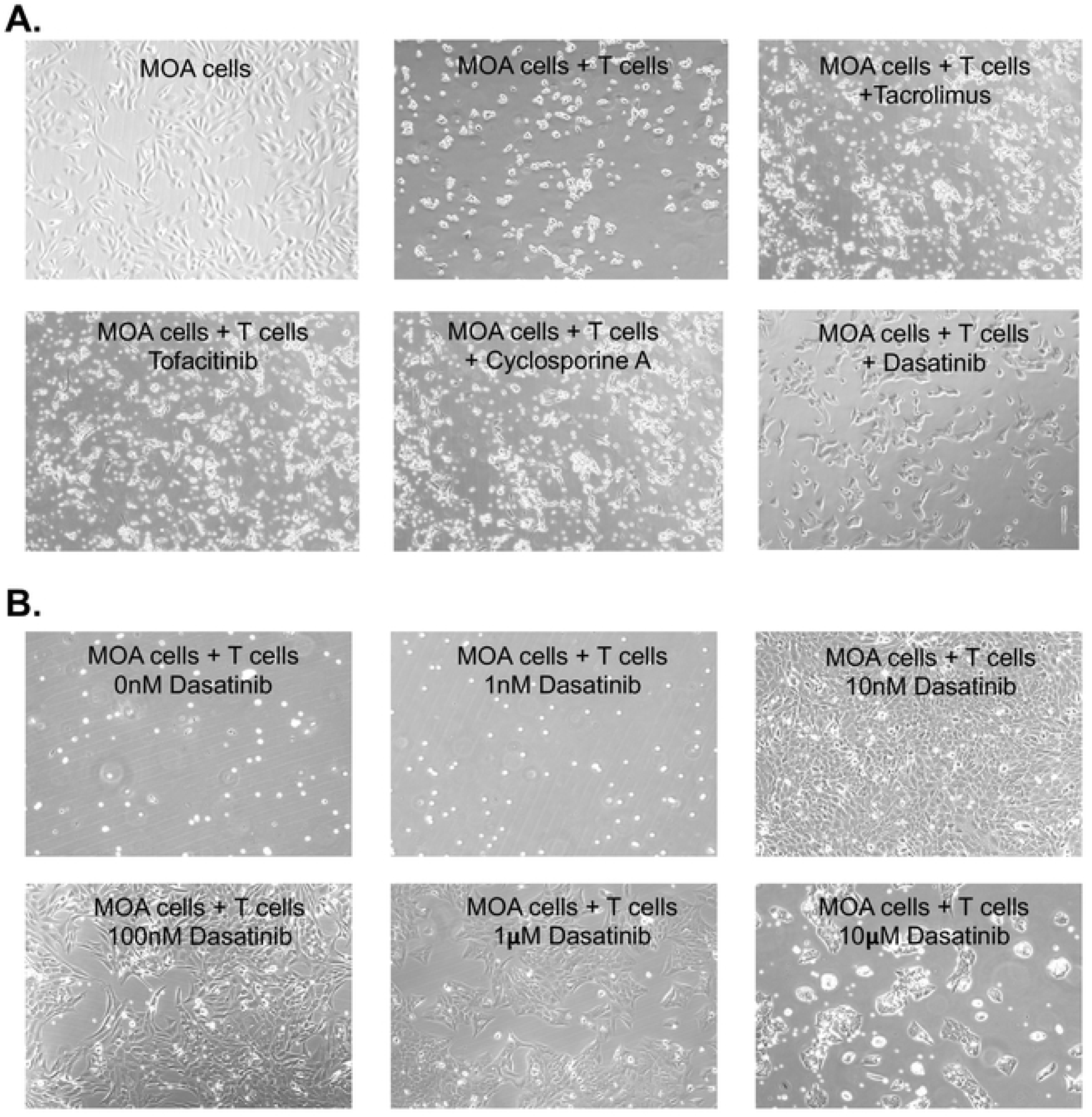
Activated T cells kill MOA reporter cells but killing is blocked by Dasatinib. **A.** MOA cells were plated at 40,000 cells per well in a 12 well plate, then cocultured with CD4 T cells that been activated with PHA/IL-2 at 500,000 cells per well. At 24h post coculture, the T cells were removed, and the remaining MOA cells were visually examined. In parallel cultures, 10μM of Tacrolimus, Tofacitinib, Cyclosporine A or Dasatinib were included. Representative images are shown. **B.** Same as A but with varying concentrations of Dasatinib included.

### Dasatinib reduces HIV infection of MOA cells but restores detection of infected cells in the presence of high density activated CD4 T cells

Although Dasatinib potently protects MOA cells from killing by activated CD4 T cells, Dasatinib has also been shown to inhibit HIV infection^27–29^. Therefore, to assess its impact on HIV infection of MOA cells, we infected MOA cells with HIV (NL4-3) in the presence of 10nM Dasatinib or control vehicle. Notably, treatment with Dasatinib significantly reduced the fraction of mCherry positive cells by ∼50% at low multiplicity of infection (MOI), indicating an antiviral effect at this concentration, although the antiviral effect was less pronounced at higher viral MOI (**Figure 6A**). This was also true for MOA cells that were cocultured with a low density of HIV infected activated CD4 T cells (**Figure 6B**). However, in the presence of high density activated, ex-vivo HIV infected CD4 T cells, the number of mCherry positive cells increased significantly in the presence of 10nM Dasatinib compared to control vehicle. The viability of MOA cells after co-culture with the ex-vivo, infected T cells was also measured, and treatment with Dasatinib significantly increased the viability of the MOA cells at high densities of activated CD4 T cells (**Figure 6C**). Taken together, these results suggest that 10nM Dasatinib does have an antiviral effect that could potentially interfere with the sensitivity of the MOA assay, but Dasatinib nevertheless increases transmission of HIV from high density activated CD4 T cells to MOA cells by preventing T-cell mediated killing of the MOA cells. Dasatinib has been previously reported to inhibit HIV infection by promoting dephosphorylation of the viral restriction factor SAMHD1^30–32^. Interestingly, when we examined the SAMHD1 expression and phosphorylation in MOA cells after 48h of 10nM Dasatinib exposure, we observed that the abundance of both total SAMHD1 and phosphor-SAMHD1 were unchanged, suggesting that a different mechanism contributes to the antiviral effect of Dasatinib in these conditions (**S5 Fig**).

**Figure 6:**
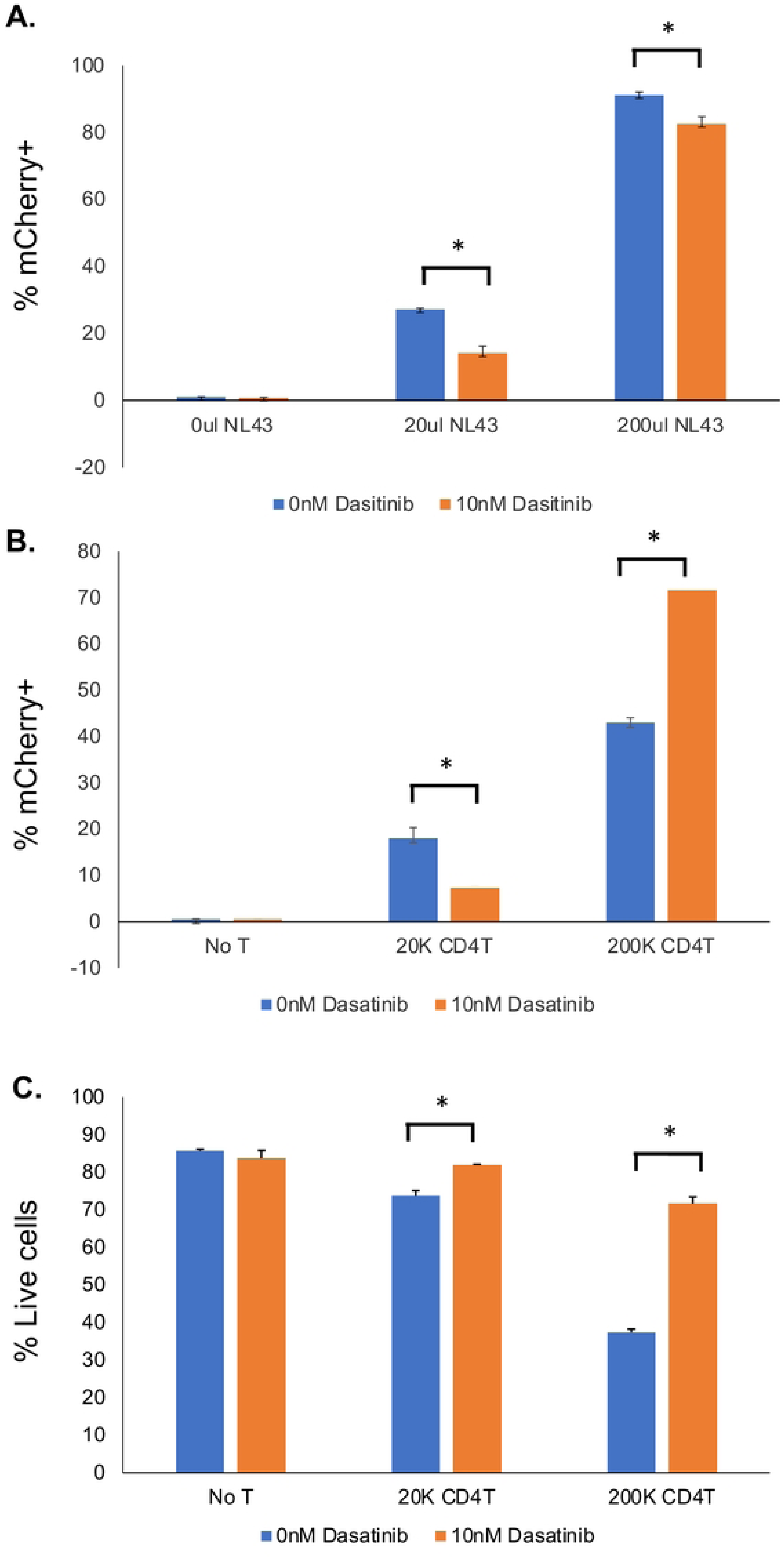
Dasatinib reduces HIV infection and transmission but improves MOA cell viability in coculture and rescues detection of infected cells. **A.** MOA cells were plated in 12 well plates at 40,000 cells per well, then infected with different quantities of HIV (NL4-3) in the presence of 10nM Dasatinib or control vehicle. At 48h post infection, the fraction of infected (mCherry positive) cells was determined by flow cytometry. **B.** MOA cells were plated in 12 well plates at 40,000 cells per well, then co-cultured with different amounts of CD4 T cells that had been activated with PHA/IL-2 then infected with NL4-3 for 48h. At 48h post coculture, the fraction of infected (mCherry positive) MOA cells was determined by flow cytometry. **C.** Viability of MOA cells from B was determined by staining with Zombie Violet (ZV) and flow cytometry. ZV negative cells were determined to be viable cells. Each bar represents an average from three replicates. Error bars represent the standard deviation of the mean. Asterisk indicates significant difference between conditions (p<0.05, Students T test)

We next tested the ability of Dasatinib to rescue MOA cell viability during high density CD4 T cell coculture in the CellRaft arrays. MOA cells were plated at four cells per raft followed by infection with HIV (NL4-3). Immediately after infection, PHA/IL-2-activated T cells were added at high density (50 cells per raft) with or without 10nM Dasitinib. In one condition, Dasatinib was washed out after 24h (**Figure 7**). When we quantified the number of mCherry+ rafts at 4dpi, we observed a potent reduction in the number of infected rafts in the presence of activated CD4 T cells, consistent with killing of the MOA cells by the CD4 T cells. Notably, addition of Dasatinib to the coculture robustly rescued detection of HIV infected MOA cells, indicated by the restoration of mCherry+ wells in the presence of Dasatinib. Washout of Dasatinib at 24h led to only a partial recovery of mCherry+ rafts, indicating that extended exposure to Dasatinib is required to protect MOA cells from killing by CD4 T cells.

**Figure 7:**
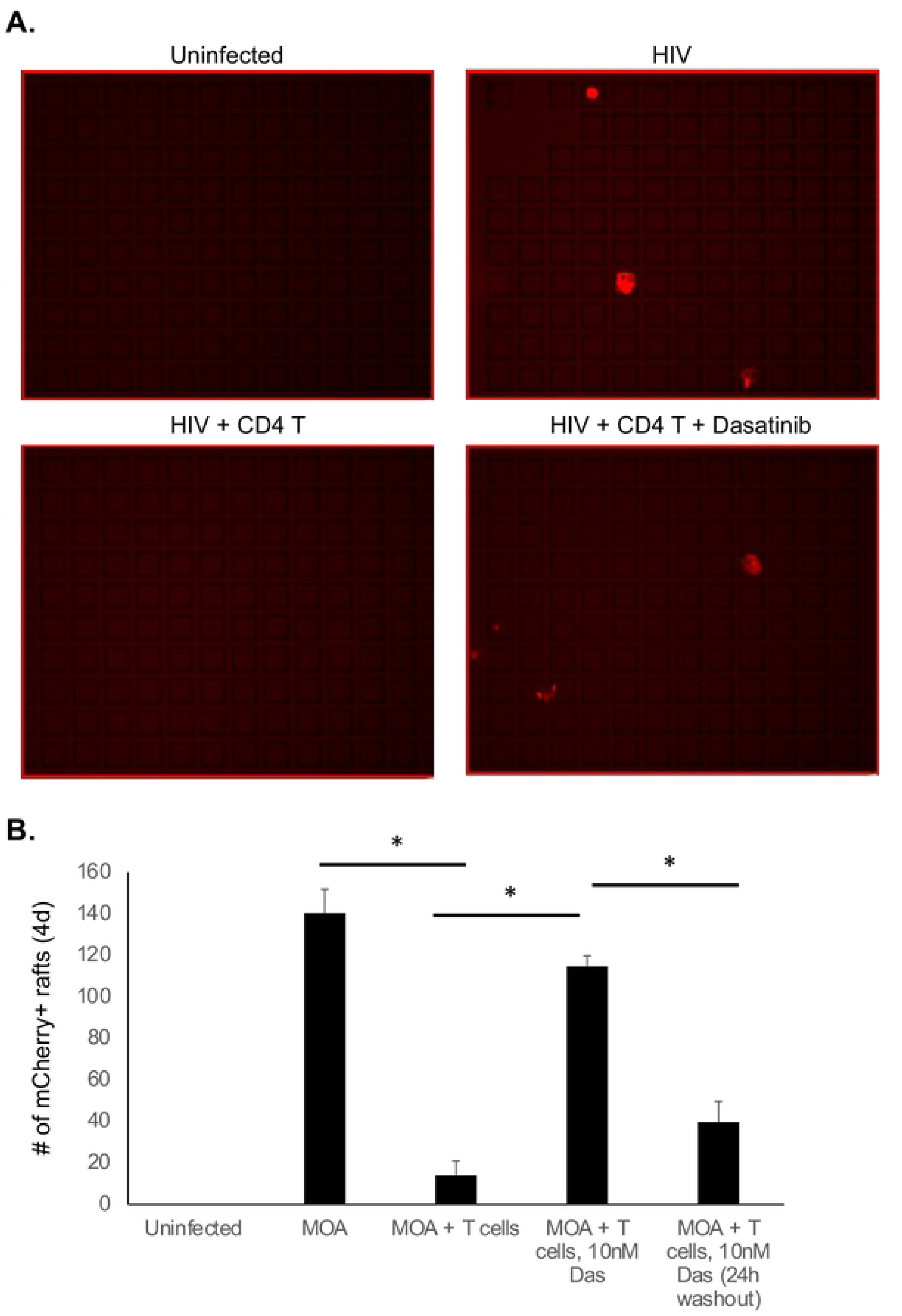
Dasatinib rescues detection of HIV infection with activated T cell coculture in CellRaft arrays. MOA cells were plated in CellRaft arrays at four cells per raft, then infected with HIV (NL4-3). At 2h post infection, media was replaced with fresh media alone or media containing PHA/IL-2 activated T cells at 50 cells per raft with or without 10nM Dasatinib (Das). In one set of arrays, the media with Dasatinib was replaced at 24h with media containing control vehicle. At day 4 of co-culture, the arrays were scanned and the quantity of infected rafts (mCherry positive) was quantified for each condition. Representative images from the arrays shown in **A.** Quantified results shown in **B.** Each bar represents the average of three replicates. Error bars represent the standard deviation of the mean. Asterisk indicates statistically significant difference between the conditions (p<0.05)

### Detection and quantification of HIV infected cells from PWH

We next examined whether we could use CellRaft arrays to detect and quantify HIV infected cells from people with HIV (PWH). To test this approach, we co-cultured peripheral blood mononuclear cells (PBMCs) from two chronically infected untreated PWH with MOA cells in CellRaft arrays for five days, then scanned the arrays and quantified the number of mCherry positive rafts. Since these samples contain actively infected cells that can be detected without additional stimulation of the cells, the PBMCs were added without prior activation and without Dasatinib. Significantly, we could detect mCherry+ rafts containing HIV outgrowth events for both donors (**Figure 8A**). We then quantified these events for each donor and calculated the number of infected cells per million PBMCs – 23/M for donor 1 and 201/M for donor 2. To confirm that this approach was accurately detecting HIV infected cells, we extracted 36 mCherry+ rafts, 12 mCherry-rafts from an infected array, and 12 rafts from an uninfected array, then performed quantitative PCR (qPCR) for HIV Gag RNA. We detected abundant Gag RNA in mCherry+ rafts but not in mCherry-rafts from the same array nor from rafts extracted from an array containing uninfected cells, validating the accuracy of the mCherry signal for identifying infected rafts (**Figure 8B**). We then further characterized viral RNA extracted from the infected rafts by sequencing the viral Gag and Envelope regions. From this we recovered sequences from individual infected cells in the original PBMC sample. Phylogenetic analysis of these sequences revealed a diverse population of infected cells as well as some cells with apparently identical sequences for Gag, while, as expected, Env sequences were more diverse (**Figure 8C**). Using this approach to recover and sequence HIV genomes from individual infection events has the potential to be useful for analysis of the genetic diversity of the reservoir.

**Figure 8:**
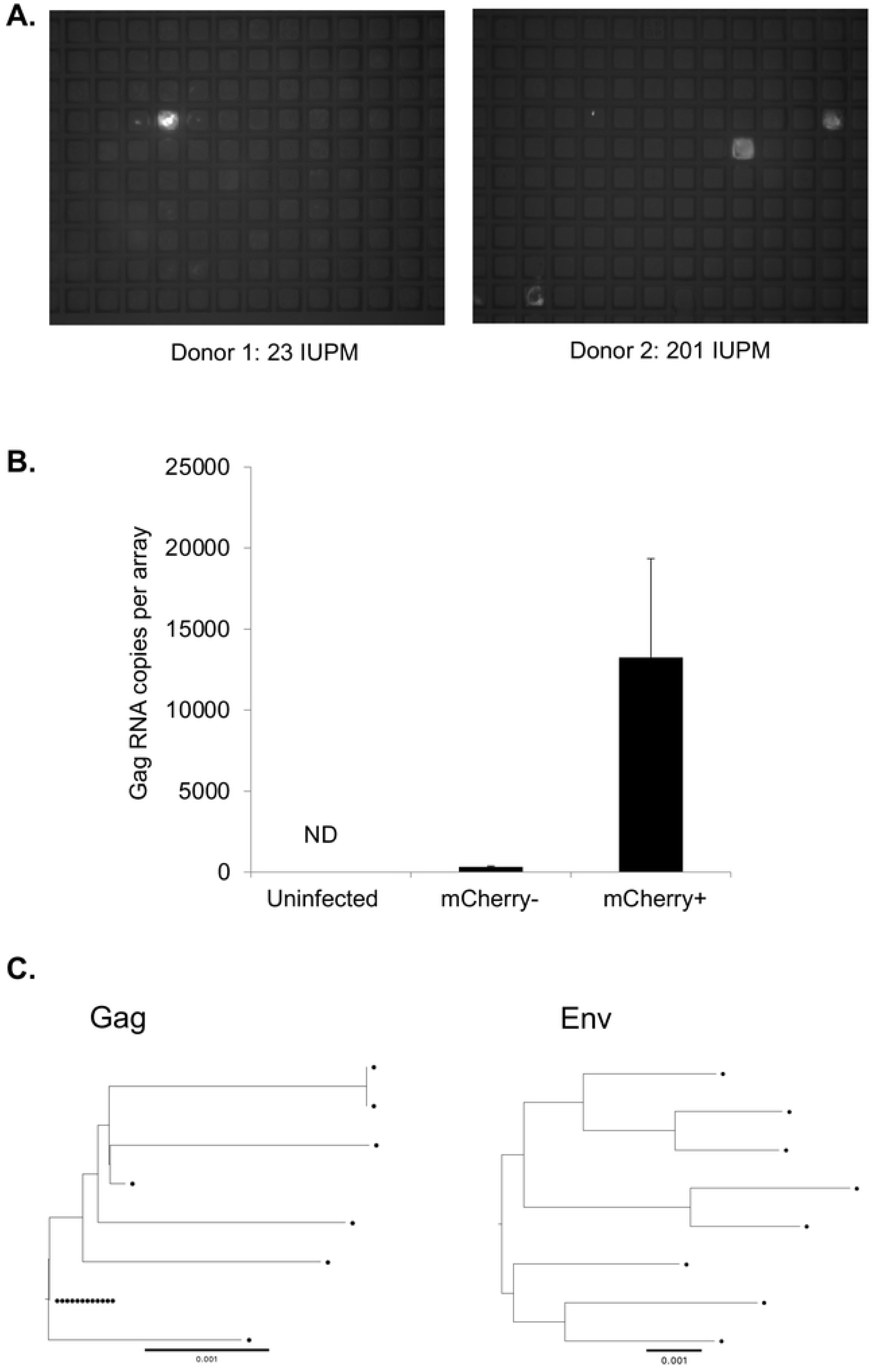
Detection of HIV infected cells from untreated PWH. **A.** MOA cells were plated in CellRaft arrays at 4 cells per raft, then overlaid with peripheral blood mononuclear cells (PBMCs) from untreated PWH at a density of 50 cells per raft. After five days of coculture, the arrays were scanned, and mCherry positive rafts were detected. **B.** 60 rafts were extracted from the arrays including mCherry positive and mCherry negative rafts and RNA was isolated from each raft. The individual rafts were then quantified for HIV Gag RNA by qPCR. **C.** Sequencing of partial Gag (HXB2 1195-1726) (left panel) and 3’ Half Genomes (HXB2 4923-9604) (right panel) RNA from virus from extracted mCherry rafts. Each sequence is represented by a circle. The reference bar indicates a phylogenetic distance of 0.001% nucleotide difference.

We then examined whether we could detect infected cells in PWH on suppressive antiretroviral therapy. This represents a more challenging goal than for untreated PWH, since infected cells are rare during therapy. To do so, MOA cells were first seeded in hexaquad CellRaft arrays with 154,000 individual rafts. PHA/IL-2 activated CD4 T-cells from PWH on ART were then added at 50 cells per raft (7.5M cells per array) along with 10nM Dasatinib to preserve MOA cell viability. Arrays were scanned daily for 7 days of co-culture and the number of mCherry+ rafts were quantified. The infectious units per million cells (IUPM) was then calculated based on the cumulative number of mCherry+ rafts over the time course of the experiment, divided by the total number of CD4 T cells plated. In all, cells from seven PWH on ART were analyzed, along with cells from four uninfected donors that were used as controls. We observed rare outgrowth events in the infected arrays, and the number of mCherry wells was significantly greater for the infected samples than for the uninfected controls, indicating that we were detecting HIV outgrowth. The infectious units per million (IUPM) from the MOA assay was compared to the previously measured QVOA IUPM estimates for the same donors (**Figure 9**). This comparison showed a linear relationship between the QVOA and MOA IUPM (r2=0.68). A significant background signal in the MOA at this scale due to rare spontaneous mCherry expression in MOA cells lead to an overestimation of the IUPM (compared to the QVOA IUPM) and will likely complicate the detection and quantification of the reservoir in PWH with small reservoirs (IUPM<1). Nevertheless, we were able to detect a clear signal above background from PWH with large reservoirs (IUPM>2). With further assay optimization this background fluorescence could potentially be minimized and a more sensitive version of this assay to support clinical trials for HIV cure may be established.

**Figure 9:**
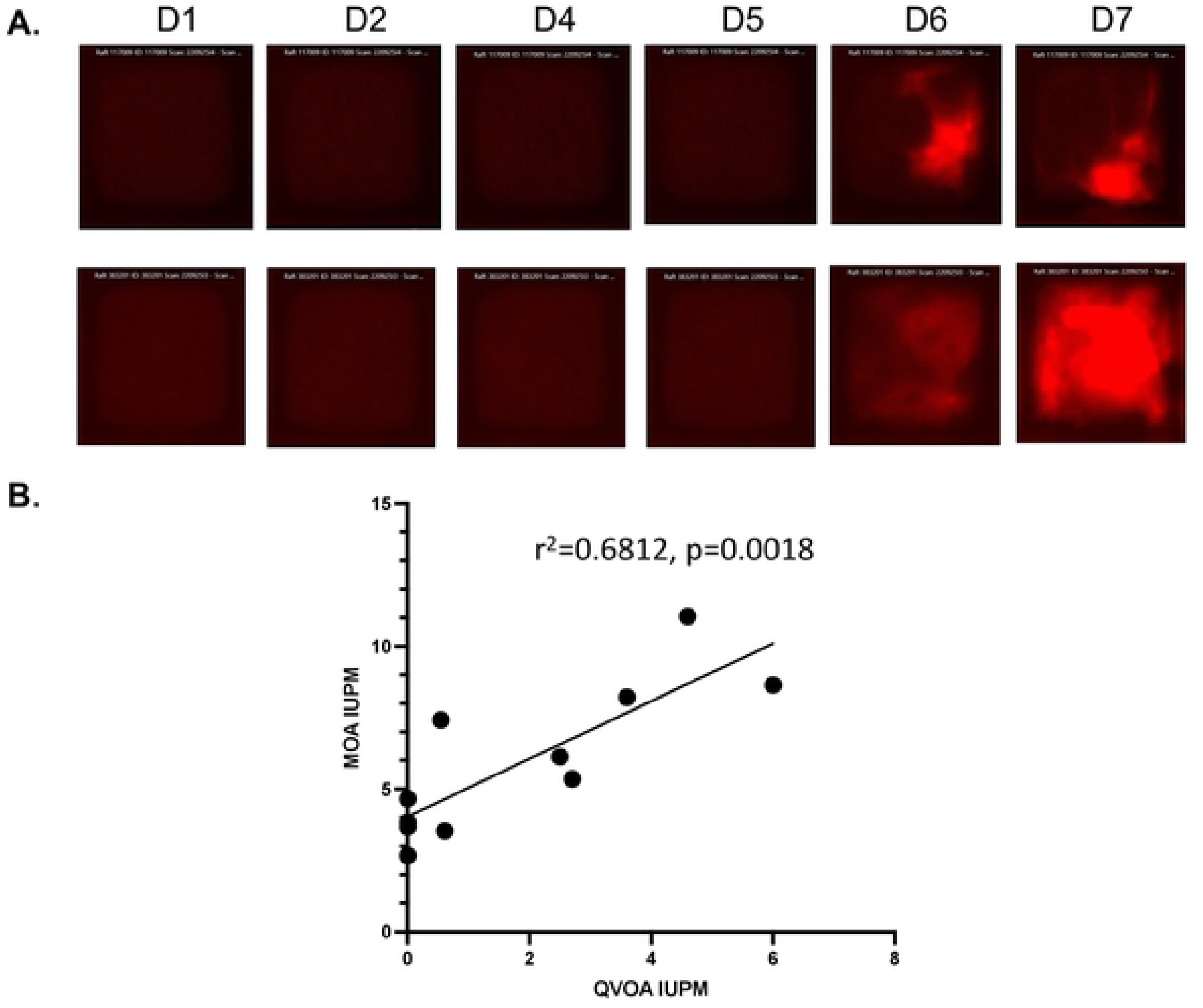
Detection of HIV infected cells in PWH on ART. Resting CD4 T cells from 7 PWH on ART and 4 serogenative controls were activated for 48h with PHA/IL-2, then washed and co-plated in hexaquad CellRaft arrays at 50 T cells per raft with MOA cells (4 cells per raft) in the presence of 10nM Dasatinib. Arrays were scanned daily for 7 days of coculture. **A.** Examples of two rafts with evidence of viral transmission are shown. **B.** Quantification of total mCherry+ rafts for each donor over the time-course calculated as infectious units per million cells (IUPM) and compared to known QVOA IUPM values for the same donors. Pearson correlation r^2^ and statistical significance are shown.

## Discussion

A key requirement for any HIV cure strategy is an ability to measure changes in the size of the viral reservoir. Several assays have been developed to accomplish this, but each of these assays comes with its own advantages or disadvantages^33,34^. The two mostly widely used assays to quantitate the HIV reservoir, IPDA and QVOA, have both provided useful insights in studies of PWH. The IPDA is rapid, high-throughput and scalable, but does not completely distinguish between defective and intact viruses and provides no information about the ability of a given provirus to reactivate and reseed infection. While QVOA, on the other hand, more precisely detects intact and replication/reactivation competent proviruses, but it underestimates the true size of the reservoir. Additionally, the assay is time-consuming, technically challenging, and not scalable to large clinical trials. As such, there is still a need for a high throughput, scalable assay for measuring the reactivation and replication competent reservoir. The microwell outgrowth assay we have developed aims to achieve this goal. While additional optimization of this approach will be required for clinical deployment, we established an assay that is capable of a comparatively quick (5-7 days) and simple quantification of infected cells from untreated PWH and ART-treated PWH with large reservoirs. Additionally, the extraction capability of the assay design permits recovery of virus sequences from individual cells in clinically relevant samples. Our hope is that this assay may be further optimized and applied to clinical trials to enable HIV cure efforts in the future.

In a direct comparison with the QVOA we observed a significant correlation between IUPM measured by MOA and IUPM measured by QVOA, indicating measurement of viral outgrowth from the latent reservoir. It is important to note, however, that this analysis depended on the use of samples for PWH with large reservoirs (IUPM>2), while quantification of the reservoir in PWH with smaller reservoirs was less successful. The MOA nevertheless offer some advantages over the QVOA. The MOA is relatively simple and quick to set up and requires no additional manipulation (e.g. addition of target cells to expand virus) after the initial plating of cells. Scanning and analysis can be carried out in an automated fashion, without the need for additional secondary assays such as p24 ELISA. As such the MOA may be more scalable to a high throughput format than the QVOA.

During the course of this study, we also observed that activated human CD4 T cells potently kill the HOS-derived MOA cells. While the precise mechanism behind this killing phenomenon is unknown, it was completely blocked by exposure to 10nM Dasatinib. Dasatinib is a tyrosine kinase inhibitor suggesting a key role for kinases in MOA cell killing. We also found that 10nM Dasatinib reduces HIV infection of MOA/HOS cells at this concentration through a mechanism that does not involve changes to the phosphorylation status of the HIV restriction factor SAMHD1. These observations may be useful for other efforts to co-culture activated human T cells with immortalized cell lines.

In its current form, the principal limitation to this assay with respect to reservoir quantification is its sensitivity. While we were able to detect and quantify rare infected cells, the background false positive rate (3-4IUPM) will make it challenging to use this assay for routine quantification of the reservoir in its current form. As such, additional work will likely be required to reduce this background signal and improve sensitivity. This could potentially be achieved through a number of approaches. Image analysis tools and machine learning could, for example, be used to more accurately distinguish false positive rafts from true infected rafts. Additional engineering of the MOA cells to eliminate any remaining HIV restriction factors that inhibit outgrowth could enhance the signal to background ratio. Further optimization of the reporter transgene or it’s integration site could also improve the signal to background ratio by reducing the number of spontaneous activation events. Finally, further optimization of the CD4 T cell stimulation conditions to reactivate a larger fraction of the reservoir could make this approach practical for quantifying the HIV reservoir.

## Acknowledgements

This work was supported by a Phase 2 SBIR award to Cell Microsystems (Durham NC) and EPB (HHS75N93020C00050).

## Author contributions

EPB conceived the study. Wet lab experiments were carried out by AF, MM and EPB. Data analyses were carried out by AF, EPB, MM and SJ. All authors contributed to writing the manuscript.

## Code and data availability

Raw data will be made publicly available without restriction upon publication.

## Methods and Materials

### CellRaft arrays

CellRaft arrays were provided by Cell Microsystems Inc (Durham, NC). Three formats of arrays were used – Cytosort arrays, with 40,000 rafts, quad arrays, with four separate arrays of 6,400 rafts, and hexaquad array with 24 separate arrays of 6,400 rafts (154,000 rafts total). To prepare arrays for cell culture, the arrays were washed three times with warm phosphate buffered saline (PBS) and left in PBS at 37°C overnight or until cells were ready for plating. To plate cells, suspensions of cells were added to each well and allowed to settle into individual rafts. Arrays were scanned on an AIR system (Cell Microsystems) with staged incubation at 37°C and 5% CO_2_. Scanning was performed in brightfield and in green and red channels (50ms and 200ms exposure times respectively). Arrays were scanned every 24h during cell culture beginning at 24h post plating. To detect mCherry positive rafts, Off The Air software (Cell Microsystems) was used in cytometric mode, with the threshold of positivity set at 5%. Rafts that were positive in the mCherry channel at the initial scan were assumed to be debris or false positive rafts and masked for the duration of the experiment. Rafts that were positive in both the red and green channels were assumed to be autoflourescent debris or dead cells and were also excluded, along with rafts where the mCherry signal was clearly not derived from cells.

### Cell Lines

HEK293 and HOS cell lines were obtained from ATCC and maintained in Dulbecco’s Modified Eagle Medium DMEM (LifeTech) containing additional 10% fetal bovine serum (FBS) and 10 U/mL Pennicilin/Streptomycin (LifeTech).

### Lentivirus generation and transduction

Lentiviruses encoding CD4, CCR5 and CXCR4 along with the selectable resistance markers neomycin, hygromycin and blasticidin respectively were purchased from Genecopeia (Rockville, MD). The LTR-mCherry reporter lentivirus was constructed by modification of the pLenti-Puro plasmid (Addgene) to remove of the RSV/LTR promoter and insertion of an intact HIV LTR-mCherry cassette. An intact HIV 3’ LTR was also inserted downstream of the SV40-puromycin resistance cassette. Lentiviruses for transduction of LTR-mCherry reporter and expression of HIV receptors (CD4, CXCR4, and CCR5) was generated by transfection of HEK293 cells using the Mirus TransIT transfection kit (Mirus). The resulting supernatant was cleared of debris by low-speed centrifugation (300g for 5 minutes) and filtration through a 0.45 micron filter. Viral supernatant was added to media on cells in culture followed by selection with the appropriate antibiotic 72 hours post transduction.

### Tat mRNA synthesis

A DNA template for Tat mRNA was created with a 5’ T7 promoter followed by the coding sequence for the Tat protein and an SV40 poly(A) signal (IDT). This sequence was amplified by PCR using the KAPA HiFi HotStart ReadyMix PCR kit and purified using a GeneJet PCR purification kit. Tat mRNA was then synthesized using the PCR amplified DNA using the MEGAscript T7 Transcription Kit (Thermo Fisher). The resulting RNA was purified using the RNEasy Plus Mini Kit (Qiagen). Purity and concentration were measured by nanodrop.

### Tat mRNA transfection

∼20,000 HOS cells transduced with the LTR-mCherry reporter were seeded into a Cytosort CellRaft array. Negative cells were identified by imaging in brightfield and red channels using the AIR system. After 24 hours, cells were transfected with Tat mRNA using the Mirus TransIT-mRNA transfection kit (Mirus). Cells that produced mCherry signal after transfection were isolated from the array and isolated as monoclonal populations into 96-well plates for expansion.

### Flow cytometry

To prepare cells for flow cytometry, MOA cells were first removed from plates by washing with PBS then trypsinization with 0.05% trypsin (Gibco). Trypsin was quenched with cell culture media, and the cells pelleted by low-speed centrifugation (300g, 5mins) before fixation with 4% paraformaldehyde. Fixed cells were re-pelleted and re-suspended in PBS with 2% fetal bovine serum and 1mM ETDA. Cells were then analyzed by flow cytometry using a Fortessa (Becton Dickson) cytometer. For analysis of HIV-HSA infected cells, the cells were first stained with an anti-HSA antibody (Biolegend) at 1:100 concentration before fixation and analysis. For analysis of CD4, CXCR4 and CCR5 expression, cells were incubated with antibodies at 1:50 dilution in PBS with 2% FBS and 1mM EDTA before washing and fixation as described above. Antibody clones used were FITC anti-CD4 (Biolegend, 317408), APC anti-CCR5 (Biolegend, 359121) and PE CXCR4 (Biolegend, 306505).

### Primary CD4 T cells

Human blood products from healthy donors were purchased from StemCell Technologies (Vancouver, Canada). Peripheral blood mononuclear cells (PBMCs) were first obtained by separation on a Ficoll gradient, followed by platelet removal via low-speed centrifugation, and removal of residual erythrocytes by resuspension in ACK lysis buffer for 5 minutes. Total CD4 T cells were then isolated using a StemCell CD4 isolation kit, and purity checked by staining with anti-CD4 and flow cytometry. Aliquots of 10-20 million CD4 T cells were frozen in 10% DMSO, 90% FBS until needed. For activation, CD4 T cells were plated at 2 million cells per mL in the presence of 1.5μg/mL phytohemagglutinin (PHA) and 60U/mL IL-2 for 48h, before washing and replating in RPMI media with 60U/mL IL-2.

### Gag RNA qPCR

Cells were sorted into TCL lysis buffer (Qiagen) before extraction of RNA using RNAClean XP beads (Beckman). RNA aliquots were the quantified for HIV Gag RNA using TaqMan Fast Virus 1-step master mix (Thermo) in an Applied Biosystems Real-Time PCR machine.

Primers and probes used were-GAGF: ATCAAGCAGCCATGCAAATGTT GAGR:CTGAAGGGTACTAGTAGTTCCTGCTATGTC GAG PROBE: FAM ACCATCAATGAGGAAGCTGCAGAATGGGA BHQ1. A standard curve consisting of a gblock covering 200 nucleotides of Gag was purchased from IDT (Coralville, IA).

### PWH Samples

PWH stably suppressed on ART (<50 HIV RNA copies/ml) for a minimum of 1 year were recruited through the University of North Carolina (UNC) Global HIV Prevention and Treatment Clinical Trials Unit and the UNC Center for AIDS Research HIV Clinical cohort to donate leukapheresis samples. The study was approved by the UNC Biomedical Institutional Review Board, and all participants provided informed consent. PBMCs were isolated from leukapheresis products by Ficoll gradient. Resting CD4 positive T cells were purified from PBMCs as previously described^35^. To estimate the frequency of resting CD4+ T cell replication-competent virus, we performed the QVOA as described elsewhere^36^. Maximum likelihood estimation statistics were used to calculate IUPMs^37^ .

### Gag Sequencing

Extracted viral RNA was used to produce a Gag amplicon using the Platinum SuperScript III One Step RT PCR System (Thermo Fisher 12-574-026). The 25μL reaction mix comprised 12.5μL 2x Buffer, 1μL 20μM F1195 5’- GTCAGCCAAAATTACCCTATAGTGC, 1μL 20μM R1726 5’-CAACAAGGTTTCTGTCATCCAATTTTTTAC, 1μL enzyme mix and 5μL template RNA. The PCR reaction conditions were 50°C for 30 min, 94°C for 2 min, 40 cycles of 94°C 15s, 50°C 30s, and 68°C 1min, followed by 68°C for 5 min. PCR products were visualized on a gel and sequenced with Sanger sequencing. Sequences were trimmed for quality and aligned to produce final sequences using Sequencher 5.4.6, compressed to unique sequences, and aligned using Muscle 5.1. The phylogenetic tree was made using Ninja v1.2.2.

### Half Genome Sequencing

Extracted RNA was converted into cDNA with an Oligo dT primer and Superscript III (Invitrogen). The 3’ half genome was then amplified with two rounds of PCR, the first using 4886F 5’-AATTCAAAATTTTCGGGTTTATTACAG and R3B3R 5’-ACTACTTGAAGCACTCAAGGCAAGCTTTATTG, followed by a nested PCR using 2.VIF1C 5’-GGGTTTATTACAGGGACAGCAGAG and Ofm19 5’-GCACTCAAGGCAAGCTTTATTGAGGCTTA, labeled with unique symmetrical Pacbio barcodes for each amplicon. The amplicons were then pooled, purified with gel extraction, and prepared into a sequencing library using the SMRTbell 3.0 Kit (Pacbio). The library was sequenced on the Pacbio Sequel IIe and the sequences were processed using SMRTlink 12.0.0 and Sequencher 3.4.1, and phylogenetic trees were constructed using Muscle 5.1 and Ninja v1.2.2.

**<u>Captions:</u>**

S1 Fig

S2 Fig

S3 Fig

S4 Fig

S5 Fig

